# HILAQ: A Novel Strategy for Newly Synthesized Protein Quantification

**DOI:** 10.1101/096925

**Authors:** Yuanhui Ma, Daniel B. McClatchy, Salim Barkallah, William W. Wood, John R. Yates

## Abstract

Here we describe a new strategy, HILAQ (<u>H</u>eavy <u>I</u>sotope <u>L</u>abeled <u>A</u>zidohomoalanine <u>Q</u>uantification), to rapidly quantify the molecular vulnerability profile to oxytosis. HILAQ quantified more than 1,900 <u>N</u>ewly <u>S</u>ynthesized <u>P</u>roteins (NSP) after 1h pulse labeling in HEK293T cell line. Compared with the previous published QuaNCAT protocol, the HILAQ strategy simplifies newly synthesized proteome analysis and achieves higher sensitivity.

## Introduction

Measuring how proteomes respond to perturbations or disease is crucial to understand the underlying mechanisms involved. Typically strategies to measure how cells respond have used techniques to measure relative gene or protein expression between perturbed and non-perturbed states and these methods have limitations in terms of scale, dynamic range and pathway efficacy. The details of biochemical mechanisms are harder to infer from expression analysis in part because of limitations associated with presumptions about co-regulation having mechanistic importance. A new method has been developed using a bioorthogonal non-canonical amino acid that is inserted into newly synthesized proteins that can then be enriched and measured. Dietrich et al synthesized azidohomoalanine (AHA) an analog of methionine that can be incorporated into proteins using the normal translational machinery and then proteins enriched with AHA can be enriched using click chemistry (1). AHA is readily incorporated into proteins, simply by adding AHA to media for cultured cells or to food pellets for animals(2), and can be used to target the most relevant proteins to a perturbation. Most analyses using AHA have been qualitative, but recently Eichelbaum et al used AHA labeling combined with metabolic stable isotope labeling to quantitate low-abundance, newly secreted proteins by mass spectrometry (MS)(3) as well as to quantitate new proteins from stimulated T-cells (QuaNCAT)(4). In the QuaNCAT method, AHA is exploited to enrich new proteins using avidin beads and heavy isotope labeled amino acids (AA) confirm proteins as newly synthesized and allow quantification in the MS. We propose a new strategy for quantification of new proteins using a new AHA molecule (**Fig.1a**), which is synthesized with heavy stable isotopes (heavy-AHA or hAHA). This strategy, HILAQ (<u>H</u>eavy <u>I</u>sotope <u>L</u>abeled <u>A</u>zidohomoalanine <u>Q</u>uantification), enables new protein enrichment, confirmation and quantification using a single stable isotope labeled molecule and employs peptide-level enrichment of hAHA.

**Figure 1.**
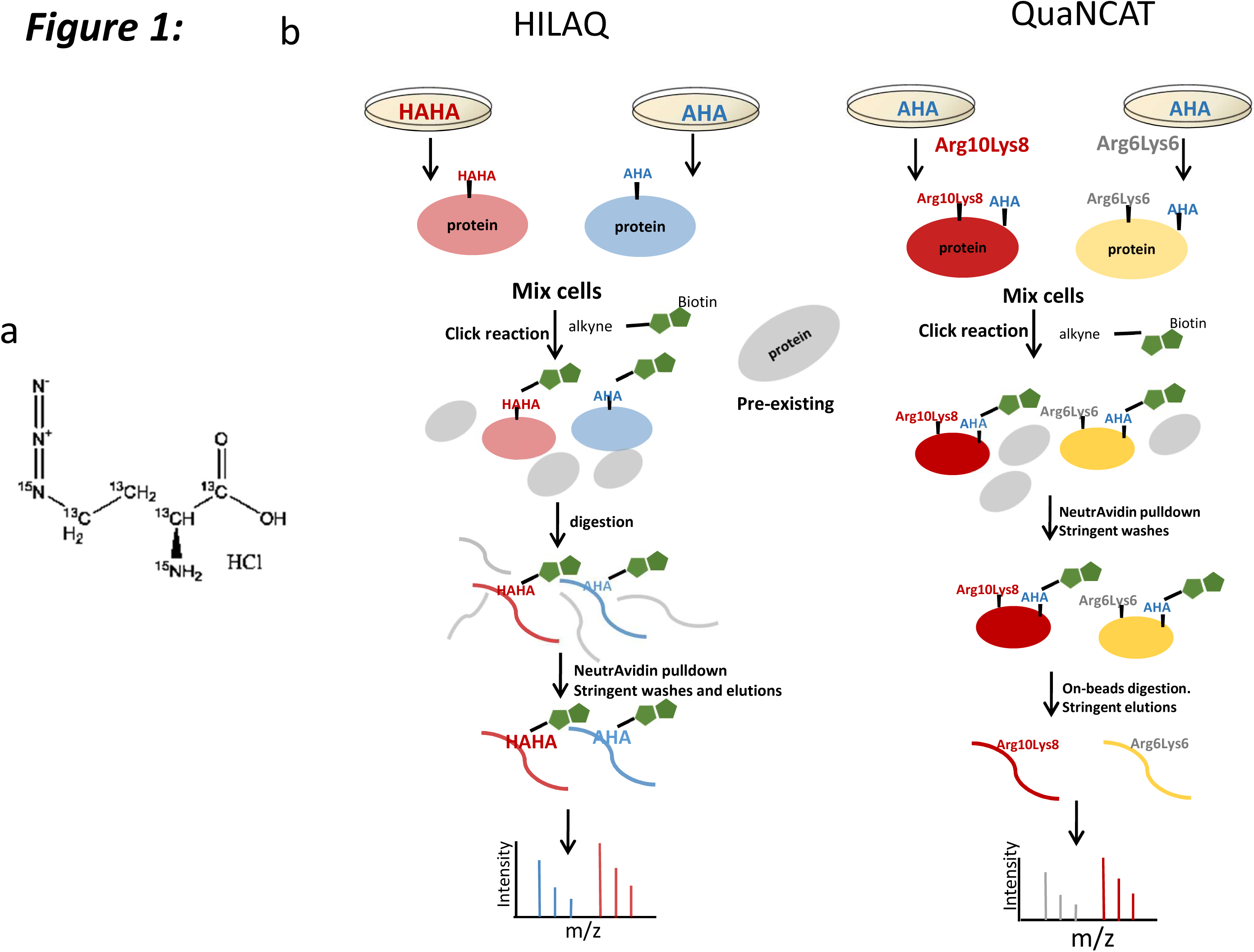
Chemical structure of heavy AHA and schematic workflows. (**a**) Two ^14^N and four ^12^C are substituted with ^15^N and ^13^C in heavy AHA. (**b**) Workflows of HILAQ and QuaNCAT. Cells are metabolically labeled with “heavy” AHA (HAHA) or “light” AHA (AHA) in the HILAQ strategy. In the QuaNCAT strategy, cells are metabolically labeled with a combination of “light” AHA, ^13^C6^15^N2-L-lysine (Lys8) and ^13^C6^15^N4-L-arginine (Arg10) or “light” AHA, ^13^C6-L-lysine (Lys6) and ^13^C6-L-arginine (Arg6). After 1h pulse labeling, plate one and two are mixed at 1:1(wt/wt) for each protocol. For both strategies, a click reaction is performed on the mixtures followed by protein precipitation. For QuaNCAT samples, AHA proteins are enriched by neutravidin beads, followed by on-bead tryptic digestion to obtain NSP peptides. The HILAQ samples are processed similarly except the neutravidin enrichment step is performed on peptides after tryptic digestion, followed by AHA peptide elution from the beads.

## Materials and Methods

### Cell preparation

HEK293T or HT22 cells were maintained in DMEM (Gibco) supplemented with 10% FBS (Invitrogen) and 1% penicillin/streptomycin (Gibco) in a humidified incubator at 37 °C and 5% CO2. Before labeling, cells were washed with warm PBS (Gibco) twice and incubated in labeling media (Hank’s Balanced Salt Solution with Glutamax, 10% dialyzed FBS, 4mM MgCl, 4mM CaCl, 1mM Sodium Pyruvate, 1% penicillin/streptomycin) for 30 min. For the HILAQ strategy, cells were incubated in labeling media with either 1mM “Heavy” AHA (^13^C_4_, ^15^N_2_, Cambridge Isotope Laboratories) (plate one) or 1mM “Light”AHA (Cambridge Isotope Laboratories) (plate two) for 1h.

For the QuaNCAT strategy, cells were incubated in labeling media with either “medium” 0.5mM Llysine(13C6, Cambridge Isotope Laboratories), 0.5mM “medium” L-Arginine (^13^C6, Cambridge Isotope Laboratories) (plate one) or 1mM “Light” AHA or “Heavy” L-lysine (^13^C6, ^15^N2, Cambridge Isotope Laboratories), “Heavy” L-Arginine (^13^C6, ^15^N4, Cambridge Isotope Laboratories) (plate two), and 1mM “Light” AHA for 1h.

For each HILAQ or QuaNCAT biological replicate, 2 plates (1 and 2) of 7×10^6^Cells were washed with warmed PBS, resuspended in TE buffer with protease and phosphatase inhibitor cocktail (Roche), then sonicated using a tip sonicator. Protein concentration was determined with a Pierce BCA protein assay (Life Technologies) prior mixing the cells 1:1(wt/wt).

### Click chemistry

For each biological replicate, 2mg of a 1:1 mixture was centrifuged. The supernatants were divided into 5 aliquots. The pellets were dissolved with 0.5% SDS in 200ul PBS, sonicated, heated on 100 °C for 10min and divided into 5 aliquots after cooling to room temperature. A click reaction was performed on each aliquot as previously published(5). Briefly, for each click reaction, the following reagents were added in this order: 1) 30 μl of 1.7mM TBTA, 2) 8 μl of 50 mM Copper Sulfate, 3) 8 μl of 5mM Biotin-Alkyne (C_21_H_35_N_3_O_6_S, Click Chemistry Tools), and 4) 8 μl of 50 mM TCEP. PBS was then added to a final volume of 400 μl and reactions were incubated for 1 hour at room temperature. All reactions were combined and TCA was added to 25%. TCA precipitation was performed overnight on shaker at 4°C.

### QuaNCAT (Biotin protein enrichment and on-beads digestion)

Precipitated proteins were dried at room temperature and resuspended in 8M Urea with 20% SDS. 150ul neutravidin agarose resin (ThermoScientific) was added and incubated for 2 h at room temperature while rotating. The resin was then washed with 1ml PBS, then PBS with 5% acetonitrile, and a final wash of PBS. The resin was centrifuged at 500 × g for 30 sec, and supernatant was removed for each wash. The resin was resuspended in 50 μl 8M Urea and 50 μl 0.2% MS compatible surfactant ProteaseMAX (Promega) in 50mM ammonium bicarbonate, then reduced, alkylated, and digested with trypsin as previously described(6). The resin digest solution was centrifuged at 500 × g for 2 min and supernatant was transferred to a new tube. Then 100 ul 50 mM ammonium bicarbonate was added to the resin and the sample was centrifuged at 500 × g for 2 min. The supernatant was removed and combined with the first supernatant. 5% formic acid was added to the combined supernatant before centrifugation at 13,000 × g for 30 min. Supernatant was transferred to a new tube, and loaded on MudPIT column.

### HILAQ (Digestion and biotin peptide enrichment)

Precipitated pellets were resuspended in 50 μl 8M Urea and 50ul 0.2% MS compatible surfactant ProteaseMAX (Promega) in 50 mM ammonium bicarbonate, then reduced, alkylated, and digested with trypsin as previously described(6). The digestion was then centrifuged at 13,000 × g for 10min. The supernatant was transferred to a new tube and the pellet was resuspended with PBS and centrifuged at 13,000 × g for 10min. Supernatants were combined and 150 μl of neutravidin agarose resin (Thermo Scientific) was added. The resin was incubated with the peptides for 2 h at room temperature while rotating, and then washed as in the QuaNCAT protocol. The peptides were eluted with two treatments of 150 μl 80% acetonitrile, 0.2% formic acid, and 0.1% TFA on shaker for 5 min at room temperature, and another two times on shaker at 70 °C. All elutions were transferred to a single new tube. Prior to MS analysis, the elution were dried with a speed-vac and dried peptides were re-solubilized in Buffer A (5% ACN, 95% water, 0.1% formic acid).

### Mass spectrometry

Soluble peptides were pressure-loaded onto a 250-μm i.d capillary with a kasil frit containing 2 cm of 10 μm Jupiter C18-A material (Phenomenex) followed by 2 cm 5 μm Partisphere strong cation exchanger (Whatman). This column was washed with Buffer A after loading. A 100 μm i.d capillary with a 5 μm pulled tip packed with 15 cm 4 μm Jupiter C18 material (Phenomenex) was attached to the loading column with a union and the entire split-column (loading column–union–analytical column) was placed in line with an Agilent 1100 quaternary HPLC (Palo Alto). The sample was analyzed using MudPIT, which is a modified 11-step separation described previously(7). The buffer solutions used were Buffer A, 80% acetonitrile/0.1% formic acid (Buffer B), and 500 mM ammonium acetate/5% acetonitrile/0.1% formic acid (Buffer C). Step 1 consisted of a 10 min gradient from 0-10% Buffer B, a 50 min gradient from 10-50% Buffer B, a 10 min gradient from 50-100% Buffer B, and 20 min 100% Buffer A. Steps2 consisted of 1 min of 100% Buffer A, 4 min of 20% Buffer C, a 5 min gradient from 0-10% Buffer B, a 80 min gradient from 10-45% Buffer B, a 10 min gradient from 45-100% Buffer B, and 10 min 100% Buffer A. Steps 3-9 had the following profile: 1 min of 100% Buffer A, 4 min of X% Buffer C, a 5 min gradient from 0-15% Buffer B, a 90 min gradient from 15-45% Buffer B, and 10 min 100% Buffer A. The Buffer C percentages (X) were 30, 40, 50, 60, 70, 80, 100% for the steps 3-9, respectively. In the final two steps, the gradient contained: 1 min of 100% buffer A, 4 min of 90% buffer C plus 10% B, a 5 min gradient from 0-10% buffer B, a 80 min gradient from 10-45% buffer B, a 10 min gradient from 45-100% buffer B, and 10 min 100% buffer A. As peptides eluted from the microcapillary column, they were electrosprayed directly into an Orbitrap Elite mass spectrometer (ThermoFisher) with the application of a distal 2.4 kV spray voltage. A cycle of one full-scan FT mass spectrum (300-1600 m/z) at 240,000 resolution followed by 20 data-dependent IT MS/MS spectra at a 35% normalized collision energy was repeated continuously throughout each step of the multidimensional separation. Application of mass spectrometer scan functions and HPLC solvent gradients were controlled by the Xcalibur data system.

### Data analysis

Both MS1 and MS2 (tandem mass spectra) were extracted from the XCalibur data system format (.RAW) into MS1 and MS2 formats using in house software (RAW_Xtractor)(8). MS/MS spectra remaining after filtering were searched with Prolucid (9) against the UniProt_Human_03-25-2014 concatenated to a decoy database in which the sequence for each entry in the original database was reversed(10). All searches were parallelized and performed on a Beowulf computer cluster consisting of 100 1.2 GHz Athlon CPUs (11). No enzyme specificity was considered for any search. The following modifications were searched for HAHA analysis: a static modification of 57.02146 on cysteine for all analyses, a differential modification of 452.2376 on methionine for AHA-biotin-alkyne or 458.2452 for HAHA-biotin-alkyne. For QuaNCAT: a static modification of 57.02146 on cysteine for all analyses, a differential modification of 6.0201 on arginine and lysine for medium label, or 8.0142 on lysine and 10.0083 on arginine for heavy label. The QuaNCAT-pep search was the same as QuaNCAT except the addition of a static modification of 452.2376 on methionine for AHA-biotin-alkyne. Prolucid results were assembled and filtered using the DTASelect (version 2.0) program(12, 13). DTASelect 2.0 uses a linear discriminant analysis to dynamically set XCorr and DeltaCN thresholds for the entire dataset to achieve a user-specified false discovery rate (FDR). In DTASelect, the modified peptides were required to be partially tryptic, less than 5ppm deviation from peptide match, and a FDR at the spectra level of 0.01. The FDRs are estimated by the program from the number and quality of spectral matches to the decoy database. For all datasets, the protein FDR was < 1% and the peptide FDR was < 0.5%. The MS data was quantified (i.e. generate heavy/light ratios) using the software, pQuant(14), which uses the DTASelect and MS1 files as the input. pQuant assigns a confidence score to each heavy/light ratio from zero to one. Zero, the highest confidence, means there is no interference signal and one means the peptide signals are almost inundated by interference signals (i.e. very noisy). For this analysis, only ratios with sigma less than or equal 0.5 were counted for statistics.

## Results and Discussion

To determine how the HILAQ protocol is different from the QUANCAT protocol, HEK293T cells were starved for 30 min in medium depleted of amino acids. Then, for each protocol, two plates of cells were pulsed with different modified AAs for 1h (**Fig. 1b**). For the QuaNCAT protocol, Plate 1 was pulsed with two “medium” SILAC AA (i.e. ^13^C_6_-R and ^13^C_6_-K) and AHA, and Plate 2 was pulsed with two “heavy” SILAC AA (i.e. ^13^C_6_^15^N_4_-R and ^13^C_6_^15^N_2_-K) and AHA. For the HILAQ protocol, the plates were pulsed with AHA (Plate 1) or HAHA (Plate 2). After mixing Plate 1 and Plate 2 1:1(wt/wt) for each protocol, 2 mg of the mixture was employed for the subsequent steps. For both strategies, a click reaction (1, 5) was performed on the mixtures, followed by protein precipitation. For QuaNCAT samples, AHA proteins were enriched by neutravidin beads, followed by on-bead tryptic digestion (6) to obtain NSP peptides. The HILAQ samples were processed similarly except the DidBIT (Direct identification of Biotin Tags) protocol was followed (6), in which the neutravidin enrichment step is performed on peptides after tryptic digestion. After AHA peptides were enriched by neutravidin beads, they were eluted with 80% acetonitrile, 0.2% formic acid, and 0.1% TFA (2). Peptides from all samples were identified using a ThermoFisher Elite Orbitrap mass spectrometer using the online peptide separation MudPIT(7). For each protocol, MS data from independent biological replicates was extracted and searched (15)(16) against a human protein database. For HILAQ analysis, mass shifts on methionine, representing either AHA-biotin-alkyne or HAHA-biotin-alkyne, were used in the database search to identify NSP. For QuaNCAT analysis, mass shifts on arginine and lysine were used in the database search to identify NSP. 2,098 NSP were confidently identified from all three biological replicates of HILAQ samples, while 363 NSP were obtained from all three biological QuaNCAT samples (**Supplementary Fig. 1**). There were 256 NSP identified by both methods (**Supplementary Fig.2**). Next, the protein samples were digested without any click reaction or avidin enrichment, then analyzed by MudPIT to determine a baseline proteome measurement. These MS datasets were searched identically without any mass shifts. The number of total peptides (i.e. HILAQ-43163 and QuaNCAT-43357) and proteins (i.e. HILAQ −5949 and QuaNCAT-5865) identified were similar for both methods. There were 5330 proteins identified in baseline of both strategies (**Supplementary Fig. 3**). We defined NSP detection sensitivity as the number of NSP over this baseline. The NSP detection sensitivity was significantly increased from 6.19% using QuaNCAT to 35.2% using HILAQ (**Fig. 2a**). In addition, after affinity purification, NSP typically comprised 13.9% of the identified proteins in QuaNCAT and increased to 79.3% in HILAQ (**Fig. 2b and supplementary Table 1**), indicating that HILAQ enrichment is more robust. Quantification of the identified NSP showed similar quantification efficiencies for the two strategies (**Supplementary Table 2**). Consistent with the large increase in NSP identifications, there were 82.7% more proteins quantified by HILAQ. (**Fig. 2c**). NSP were quantified by an average of 8.78 peptides by the HILAQ protocol compared with 3.40 peptides by QuaNCAT (**supplementary Table 1**). The Gaussian distribution pattern of protein ratios and (**Fig. 2d**) the dispersion of protein ratios for the HILAQ and QuaNCAT strategies were similar (**Fig. 2e**), suggesting that HILAQ and QuaNCAT have equal accuracy. Pathway analysis was performed on the quantified proteins from the HILAQ or QuaNCAT methods. **Fig. 2f** shows that proteins quantified by HILAQ were enriched in 133 pathways. This is twice as many as with QuaNCAT, and suggests that HILAQ is a more informative strategy (**supplementary Fig. 4**). Besides using different heavy molecules, the QuaNCAT protocol employs AHA protein enrichment while the HILAQ protocol employs AHA peptide enrichment. Peptide enrichment has been demonstrated to be more sensitive than protein enrichment in biotin labeled protein identification (6). To illustrate the benefit of peptide enrichment for quantification of NSP, we developed QuaNCAT-pep, a modified QuaNCAT protocol which includes peptide enrichment rather than protein enrichment. (**Supplementary Fig. 5**). QuaNCAT-pep confidently identified twice as many NSPs than QuaNCAT (**Supplementary Fig. 6**) and NSP detection sensitivity was increased to 23.3% in QuaNCAT-pep from 6.19% in QuaNCAT (**Supplementary Fig. 7**). This analysis suggests that the improved performance of the HILAQ protocol over the QuaNCAT protocol is primarily due to their different AHA enrichment strategies rather than the use of different heavy molecules.

**Figure 2.**
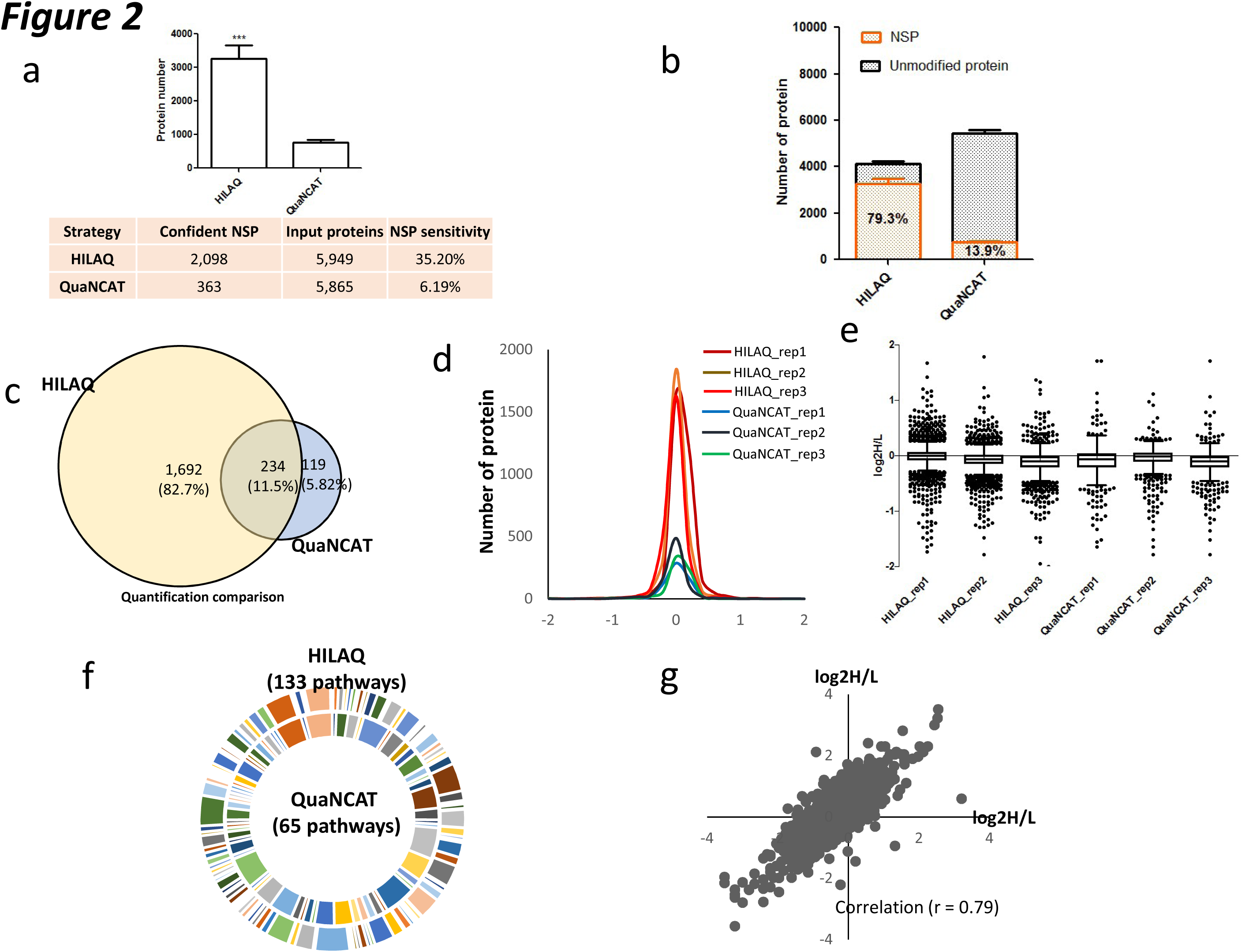
Performance of HILAQ. (**a**) The number of NSP identified by HILAQ protocol was significantly increased over QuaNCAT. In the bottom table, confident NSP means the NSP overlapped in all three biological replicates. NSP sensitivity is the ratio of modified peptide/protein identifications over baseline. (**p<0.001; one tailed unpaired t-test, comparison with QuaNCAT) (**b**) Composition of identified protein by HILAQ and QuaNCAT. (**c**) Venn diagram analysis on quantification by HILAQ or QuaNCAT. Only proteins quantified confidently by all three replicates of HILAQ or QuaNCAT were counted and compared for this analysis. (**d**) Normal distribution of replicates of HILAQ and QuaNCAT. All datasets were not normalized. (**e**) Boxplot analysis on every replicate of HILAQ and QuaNCAT using Graphpad Prism (version 5.01). The boxplot was created with Turkey whiskers. (**f**) Proteins quantified by all three replicate of HILAQ or QuaNCAT were annotated using web based tool PANTHER (http://pantherdb.org/). There are two times more pathways enriched in HILAQ than QuaNCAT. (**g**) The quantification of NSP between HT22 and HEK293Tcell lines. The peptides ratios from two independent HILAQ experiments were compared. The AHA and HAHA labels were swapped in the two experiments.

To demonstrate the usefulness of HILAQ, an in vitro model of oxytosis was analyzed. Glutamate is a major neurotransmitter in the brain but excessive extracellular glutamate can result in oxidative stress leading to cell death. This glutamate toxicity has been implicated in many neurological diseases, including cerebral ischemia, Huntington’s disease, Alzheimer’s disease (17), and amyotrophic lateral sclerosis (18). Glutamate induces toxicity through two mechanisms: excitotoxicity or oxytosis. In excitotoxicity, excess extracellular glutamate overstimulates ionotropic glutamate receptors leading to cell death initiated by an influx of calcium ions (19, 20). In oxytosis or non-receptor oxidative glutamate toxicity, excess glutamate inhibits a cystine/glutamate antiporter preventing cystine import which is the rate limiting step of glutathionine (GSH) synthesis. GSH is one of the cell’s important antioxidants to neutralize reactive oxygen species (ROS). Oxidative modification of proteins, lipids and DNA by ROS produces oxidative stress. The ROS triggers downstream signaling pathways generating increased intracellular calcium leading to cell death (21). This particular cell death is a novel form of programmed cell death with characteristics of both necrosis and apoptosis, which has been described as necroptosis (22). Increased oxidative stress and depletion of GSH is associated with aging and neurodegenerative diseases, such as Alzheimer’s Disease (23, 24). Thus, studying oxytosis may hold potential for identifying treatments for complex neurodegenerative disorders (25). The immortalized hippocampal cell line, HT22, is an in vitro model of oxytosis due to its lack of ionotropic glutamate receptors and high sensitivity to extracellular glutamate(26). It has been used as an important research tool to study the molecular details of oxytosis and investigate neuroprotective pathways. For example, HT22 cells selected for resistance to oxytosis were discovered to be less vulnerable to amyloid-beta toxicity, which is hypothesized to contribute to the pathogenesis of Alzheimer’s disease (27). HILAQ analysis was employed to compare HT22 cells to HEK293T cells to determine the molecular vulnerability profile to oxytosis. HEK293T cells were pulsed with AHA and HT22 cells were pulsed with hAHA. In the second experiment labeling of the cells was reversed. A comparison of the swap experiments between two biological replicates showed sufficient reproducibility with a Pearson correlation coefficient of 0.79 (**Fig. 2g**). In total, 818 NSPs were quantified in the swap experiments. Pathway analysis was performed on 226 proteins that were observed to have at least a twofold change between the cell types with 136 upregulated and 90 down-regulated in the HT22 cells. The most significantly (p value = 1.16 e −16) enriched pathway was cell death with 108 proteins annotated to this pathway. Cell fate is dictated by the relative activity of a complex network pro- and anti-death proteins. In accordance, 68 of these proteins have been reported decrease cell death suggesting have a protective function while 27 have been reported to increase cell death suggesting pro-death function or required for cell death to proceed (**Fig. 3**). The remaining thirteen proteins were reported to be involved in cell death without direct evidence for halting or activating the process. Many of these altered proteins have previously been connected to cell death in brain tissue and disease models. For example, BIRC6 (Baculoviral IAP repeat-containing protein 6) is part of a protein family that inhibits cell death through the inactivation of caspases and was up-regulated in HT22 cells. It has been reported that the hippocampus downregulated this protein in response to glutamate toxicity(28). In order for cell death to proceed, cells degrade BIRC6 through the proteasome (29). Interestingly, it has been reported that proteasome inhibitors can prevent oxytosis in HT22 cells (30, 31). TRAP1 (Tumor necrosis factor type 1 receptor-associated protein) is mitochondrial molecular chaperone and was down-regulated in HT22 cells. It has been reported that PINK1 (PTEN-induced putative kinase protein 1) phosphorylates TRAP1, which is essential for PINK1 to protect cells from cell death induced by oxidative stress (32). Mutations in PINK1 can cause Parkinson’s disease(PD), which is characterized by neurodegeneration of dopamine neurons induced by oxidative stress(33, 34). The major source of ROS in oxytosis is generated from the mitochondrial complex I and complex I dysfunction is implicated in PD pathogenesis (35, 36). The core subunit of complex I, NDUFV2, was up-regulated in HT22 cells. Finally, p62/SQSTM1 (Sequestosome-1) was up-regulated in HT22 cells. This protein has been localized to protein aggregates that accumulate in multiple neurodegenerative diseases (37–39). It is an autophagy receptor that binds poly-ubiquitinated proteins to direct them to lysosomes for degradation. It was reported that SQSTM1 can protect cells from cell death induced by mutant huntingtin, which causes the neurodegenerative disease, Huntington’s Disease (40). In summary, this HILAQ analysis demonstrates the validity of HT22 cells as a tool to study the molecular details of cell death involved in neurodegenerative diseases.

**Figure 3.**
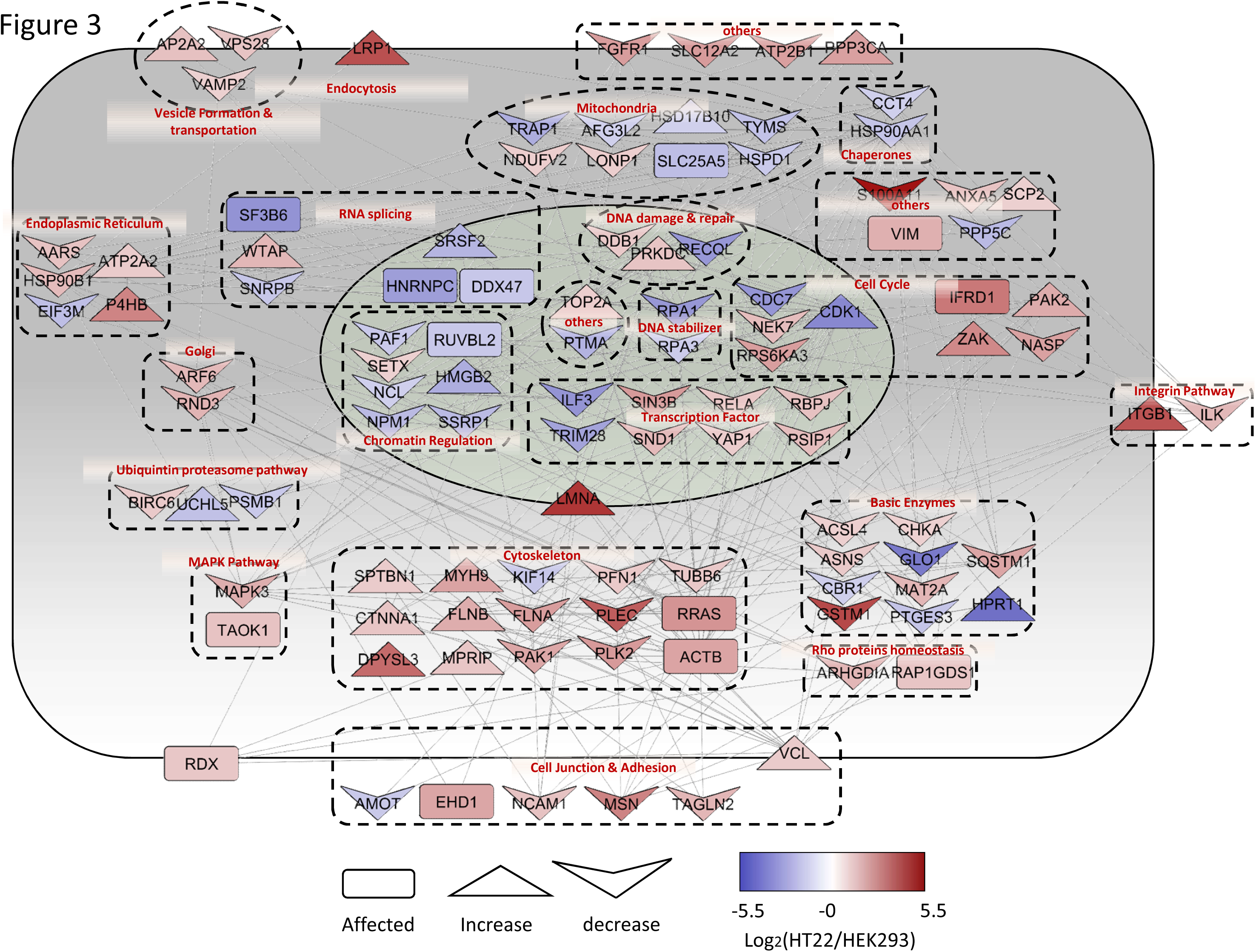
Interaction network analysis on 2 fold express changed protein after 1 h labeling in HEK293T and HT22 cells. The protein network and localization were analyzed by String and visualized by Cytoscape. The log ratio used in visualization was the average ratio of replicates. The red color represents up-regulation while the blue color indicates decrease. Different shapes used in the figure represents the effects of the protein on cell apoptosis according to previous reports.

## Conclusion

We have described a novel strategy, HILAQ, to quantify the newly synthesized proteome. Since the HILAQ strategy employs two modified AA while the QuaNCAT method employs five modified AA, HILIQ simplifies the workflow and bioinformatics analysis. While QuaNCAT employs AHA solely for enrichment of NSP and relies on the SILAC AA for confirmation and quantification of a NSP, the HILAQ method exploits the AHA molecule for enrichment, confirmation and quantification of the NSP. We also demonstrated that the quantification of NSP is greatly improved when peptide AHA enrichment is employed rather than protein AHA enrichment. Overall, the high sensitivity of this approach allows it to be used to monitor proteomic dynamics using short-time pulse labeling instead of traditional wholeproteome labeling.

## Acknowledgements

Funding from the following National Institute of Health grants: P41 GM103533, R01 MH067880, R01 MH100175 to Yates laboratory.

